# Meta-analysis of Reward Processing in Major Depressive Disorder Reveals Distinct Abnormalities within the Reward Circuit

**DOI:** 10.1101/332981

**Authors:** Tommy H. Ng, Lauren B. Alloy, David V. Smith

## Abstract

Many neuroimaging studies have investigated reward processing dysfunction in major depressive disorder. These studies have led to the common idea that major depressive disorder is associated with blunted responses within the reward circuit, particularly in the ventral striatum. Yet, the link between major depressive disorder and reward-related responses in other regions remains inconclusive, thus limiting our understanding of the pathophysiology of major depressive disorder. To address this issue, we performed a coordinate-based meta-analysis of 41 whole-brain neuroimaging studies encompassing reward-related responses from a total of 794 patients with major depressive disorder and 803 healthy controls. Our findings argue against the common idea that major depressive disorder is primarily linked to deficits within the reward system. Instead, our results demonstrate that major depressive disorder is associated with opposing abnormalities in the reward circuit: hypo-responses in the ventral striatum and hyper-responses in the orbitofrontal cortex. The current findings suggest that dysregulated corticostriatal connectivity may underlie reward-processing abnormalities in major depressive disorder, providing an empirical foundation for a more refined understanding of abnormalities in the reward circuitry in major depressive disorder.

## Introduction

Depression is a prevalent mental disorder ranked as the leading non-fatal cause of disability by the World Health Organization.^1^ Therefore, it is of paramount importance to understand its underlying neurobiological mechanisms. Over the past decade, theorists have proposed that anhedonia, one of the core symptoms of depression, is linked to reward processing dysfunction.^2–9^ In particular, many neuroimaging studies have reported that relative to healthy controls (HC), individuals with major depressive disorder (MDD) exhibit reduced activity in the ventral striatum (VS) in response to reward.^10–14^

The striatum, which can be divided into dorsal and ventral sections, is the primary input zone for basal ganglia.^16,17^ It receives afferent projections from the midbrain, amygdala, and prefrontal cortex (PFC), such as the orbitofrontal cortex (OFC), dorsolateral prefrontal cortex (dlPFC), ventromedial prefrontal cortex (vmPFC), and anterior cingulate cortex (ACC).^16,17^ It also projects to such regions as the ventral pallidum, ventral tegmental area, and substantia nigra.^17^ Many of the regions linked to the striatum, particularly prefrontal regions, have been associated with the computation and representation of reward value,^18–24^ as well as the regulation of affect and reward-related behavior in animals and healthy individuals.^25–29^ The striatum also has been proposed to play an important role in the onset and course of MDD, with longitudinal studies demonstrating that blunted activation in the VS during reward anticipation predicts the emergence of depressive symptoms and disorder^30,31^ and deep-brain stimulation studies using it as a treatment target for treatment-resistant depression.^32^

Although blunted striatal response to reward in MDD is a well-established finding in the literature,^9,33–36^ it is less clear how other regions, particularly the PFC, also may contribute to reward processing deficits in MDD. For instance, some studies have found that relative to HC, MDD exhibited greater activation in the OFC,^13,37^ dlPFC,^12,38,39^ vmPFC,^40,41^ACC,^42,43^ middle frontal gyrus,^41,43^ inferior frontal gyrus,^42,44^ subgenual cingulate,^40,44^ and dorsomedial prefrontal cortex^41^ during the processing of rewarding stimuli. In contrast, other studies have reported less activity in MDD in response to reward in the OFC,^37,43^ ACC,^12,13,37,44^ middle frontal gyrus,^13,42,44^ and frontal pole.^43^ The inconsistencies may be due to a number of factors, such as limited statistical power^45–47^ and susceptibility artifacts in the ventral portions of the PFC.^25,48,49^ Therefore, the association between prefrontal regions and MDD remains equivocal, both in terms of the *direction* (i.e., hyper- or hypo-responses) and the *location* (e.g., OFC, dlPFC, vmPFC and/or ACC) of the effect.

Inconsistencies in the literature have prompted researchers to conduct coordinate-based meta-analyses to identify common activation patterns implicated in MDD during reward processing. Although prior meta-analytic efforts^34,35,50^ have shown some overlapping findings in the striatum in MDD, there is a striking degree of anatomical disagreement across these efforts, with non-overlapping findings all throughout the brain (see Supplementary Table 1 and Supplementary Figure 1 for a complete comparison of findings across meta-analyses). The lack of agreement across meta-analyses can be due to methodological issues, such as lenient thresholding, overlapping samples, software issues,^51^ different inclusion criteria, and inclusion of region-of-interest (ROI) coordinates, as detailed in a previous review.^52^ For example, two previous meta-analyses^34,35^ corrected for multiple comparisons using the false discovery rate (FDR) approach, which has been shown to be inadequate in controlling the false positives among clusters in neuroimaging meta-analyses^53,54^ and might have contributed to the lack of agreement across studies.

**Table 1.** Peak Coordinates of Group Differences in Neural Responses to Reward

| Contrast | Cluster Size<br>(mm<sup>3</sup>) | Probabilistic Anatomical<br>Label | x | y | z |
| --- | --- | --- | --- | --- | --- |
| MDD > HC | 912 | Frontal Orbital Cortex<br>(26%), Frontal Pole (13%) | 20 | 32 | -12 |
| HC > MDD | 1768 | Subcallosal Cortex (14%) | -2 | 8 | -4 |
|  |  | Caudate (32.1%),<br>Accumbens (11.1%) | 8 | 6 | -2 |

Abbreviations: MDD, major depressive disorder; HC, healthy controls. Coordinates are x,y,z
values of the locations of the maximum activation likelihood estimation (ALE) values in
MNI space. Probabilistic labels reflect the probability that a coordinate belongs to a given
region derived from the Harvard-Oxford probabilistic atlas. For clarity, we only report labels
whose likelihood exceeds 5%.

**Figure 1.**
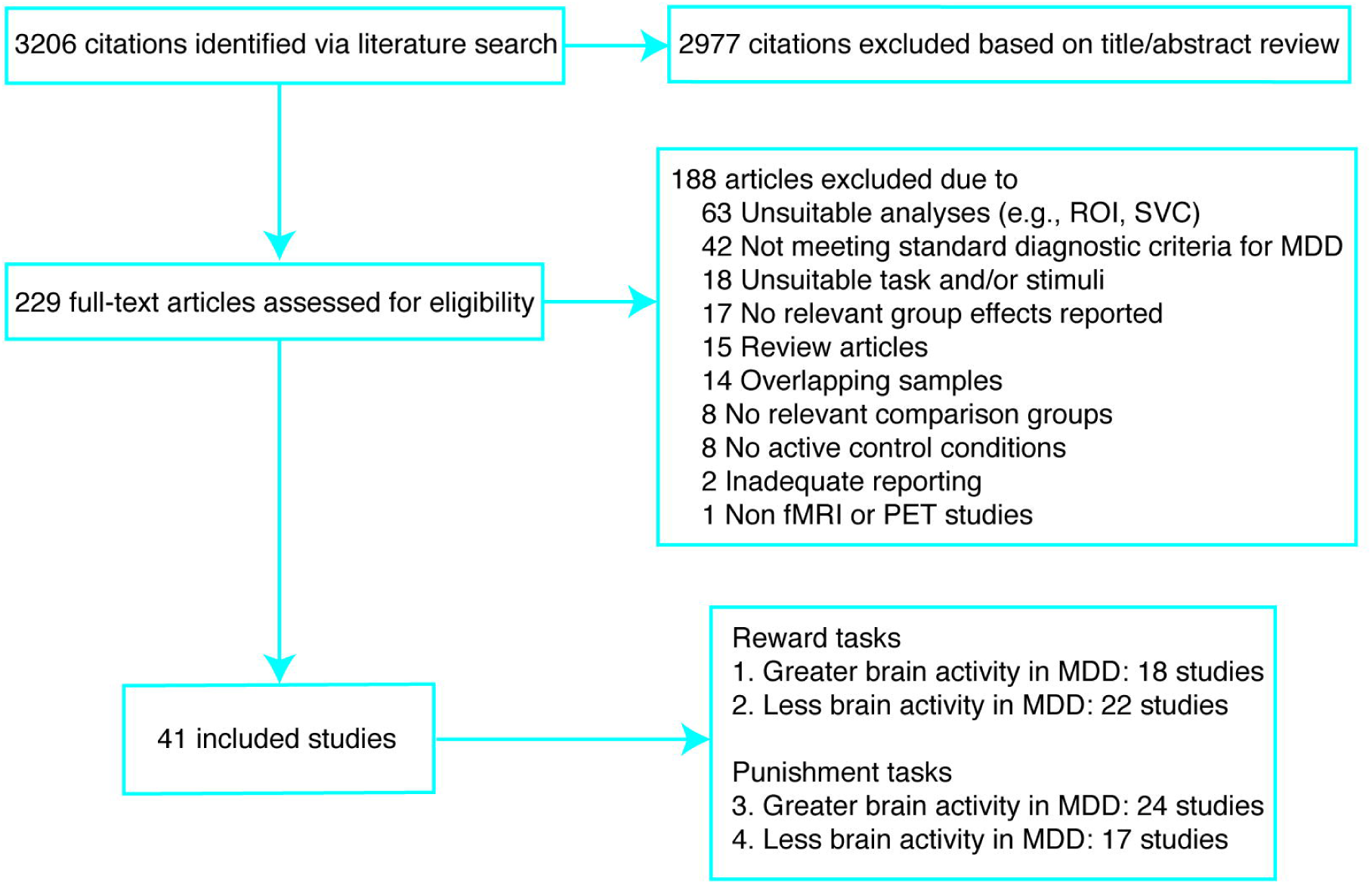
Flowchart of study selection. The systematic literature search identified a total of 41 neuroimaging studies that met the inclusion criteria, yielding 4 coordinate-based meta-analyses with at least 17 independent studies; ROI, region of interest; SVC, small volume correction; MDD, major depressive disorder.

To address these issues and extend extant work, we performed a coordinate-based meta-analysis following procedures recommended by new guidelines.^52,55^ The current work differed from previous meta-analyses on reward processing in MDD in various aspects, such as only including whole-brain studies to avoid localization bias; only including studies that used an active control condition to isolate reward-related processes; only including independent samples to avoid double counting the same participants; using more stringent thresholding criteria; having the most up-to-date literature search; and only conducting a meta-analysis when there were at least 17 eligible experiments to restrict excessive contribution of any particular studies to cluster-level thresholding.^56^

Our primary hypothesis was that the literature would consistently show that compared with HC, individuals with MDD would exhibit blunted activation of the striatum and abnormal activation of the prefrontal regions (e.g., the OFC) during the processing of rewarding stimuli. We also explored whether there were consistent neural responses to punishing stimuli in MDD relative to HC. To examine these questions, we conducted four separate coordinate-based meta-analyses testing spatial convergence of neuroimaging findings for the following four contrasts: 1) positive valence (reward > punishment or neutral stimuli; neutral stimuli > punishment) for MDD > HC; 2) negative valence (punishment > reward or neutral stimuli; neutral stimuli > reward) for MDD > HC; 3) positive valence for HC > MDD; and 4) negative valence for HC > MDD. Neutral stimuli were considered rewarding relative to punishing stimuli but punishing relative to rewarding stimuli, in view of the idea that brain activation during reward processing reflects the valence of a stimulus and follows a linear pattern.^57,58^

The comprehensive and rigorous nature of the current meta-analysis allowed us to investigate whether a quantitative synthesis of neuroimaging studies on reward-related processing in MDD would unveil common activation patterns that may be difficult to discern by individual studies due to inconsistent findings. We aimed to address two main questions. First, which brain regions show consistent hypo-responses to reward-relevant stimuli in MDD relative to HC? Second, which brain regions show consistent hyper-responses to reward-relevant stimuli in MDD relative to HC?

## Materials and Methods

### Study Selection

The current coordinate-based meta-analysis primarily followed the guidelines for meta-analyses, whenever applicable.^55,59^ We conducted a systematic literature search to identify neuroimaging studies on reward processing abnormalities in mood disorders. Potentially eligible studies published between 1/1/1997 and 8/7/2018 were identified by searching the MEDLINE, EMBASE, PsycINFO, PsycARTICLES, Scopus, and Web of Science using the grouped terms (fMRI* or PET*) AND (depress* OR bipolar* OR mania* OR manic* OR hypomania* OR hypomanic*) AND (reward* OR effort* OR decision* OR reinforce* OR habit* OR discounting* OR “prediction error” OR “delayed gratification” OR “approach motivation” OR “positive valence systems”). To enhance search sensitivity, the reference lists of the retrieved articles and review papers were further checked to identify potentially relevant articles. Although our initial goal was to investigate reward processing dysfunction in both MDD and bipolar disorder, the current meta-analysis only focused on MDD due to an inadequate number of studies on bipolar disorder (the search identified 23 studies on bipolar disorder across positive and negative valence contrasts, yielding fewer than 17 experiments for each targeted meta-analysis).

### Inclusion Criteria

We included studies that (a) used a reward and/or punishment task, (b) reported comparisons between people with MDD and HC, (c) used standardized diagnostic criteria (e.g., *DSM-IV, DSM-IV-TR*, ICD-10) to determine psychiatric diagnoses, (d) used fMRI or PET in conjunction with parametric analysis or subtraction methodology contrasting an experimental condition and an active control condition (e.g., a punishment condition, a lower-intensity reward condition, or a neutral condition) to isolate reward-related processes and identify foci of task-related neural changes, (e) reported significant results of whole-brain group analyses without small volume corrections (SVC), as non-whole-brain coordinates (e.g., ROI-based coordinates) and analyses involving SVC have been argued to bias coordinate-based meta-analyses,^55,56^ (f) reported coordinates in a standard stereotactic space [Talairach or Montreal Neurological Institute (MNI) space], and (g) used independent samples. Authors were contacted if the required information were unavailable in the published reports.

The study with the largest sample size was included if there was sample overlap between studies. Reward tasks were operationalized as involving presentation of a rewarding stimulus (e.g., winning money, favorite music, positive faces), whereas punishment tasks were operationalized as involving presentation of a punishing stimulus (e.g., losing money, negative faces). The stimuli used in the included studies of the meta-analysis reflect a reward-punishment or positive-negative continuum. For example, positive faces are considered as rewards based on previous research showing that positive faces activate the reward circuitry, that they are discounted as a function of time, that they are tradable for other rewards (e.g., money), that they reinforce work, and that people are willing to work to view positive faces and exert more effort for more positive faces.^60,61^

### Coordinate-Based Meta-Analysis

Given the inconsistency of findings in the literature of reward processing abnormalities in MDD, we used a coordinate-based meta-analytic approach and activation likelihood estimation^54,62^ to examine whether we could identify consistent activation patterns across studies. Our main analyses focused on which brain regions show consistent hypo- or hyper-responses to reward-relevant stimuli in MDD relative to HC. Coordinate-based meta-analyses were performed using GingerALE 2.3.6 (http://brainmap.org), which employs the activation likelihood estimation (ALE) method.^54,63^ The ALE method aims to identify regions showing spatial convergence between experiments and tests against the null hypothesis that the foci of experiments are uniformly and randomly distributed across the brain.^54^ It treats foci from individual experiments as centers for 3D Gaussian probability distributions representing spatial uncertainty. The width of these distributions was determined based on between-subject and between-template variability.^62^ The ALE algorithm weighs the between-subject variability by the number of participants for each study, based on the idea that experiments of larger sample sizes are more likely to reliably report true activation effects. Therefore, experiments with larger sample sizes are modeled by smaller Gaussian distributions, resulting in a stronger influence on ALE scores, which indicate the probability that at least one true peak activation lies in the voxel across the population of all possible studies.^62^ As a result, studies with larger sample sizes would be weighed more heavily relative to studies with smaller sample sizes.

The ALE method is implemented in the following steps. First, for each included study, a map of the activation likelihood is computed. Second, the maps are aggregated to compute the ALE score for each voxel. Finally, a permutation test is employed to identify voxels in which the ALE statistic is larger than expected by chance.^54,62,63^ The ALE method takes into account heterogeneity in spatial uncertainty across studies^54,62,63^ and differences in number of peak coordinates reported per cluster.^63^ This approach allows random-effects estimates of ALE, increasing generalizability of the results.^62^

Coordinate-based meta-analyses represent a departure from traditional meta-analyses.^55^ Specifically, whereas traditional meta-analyses aim to calculate pooled effect sizes to determine the direction and magnitude of an effect based on a body of literature, coordinate-based meta-analyses evaluate whether the location of an effect is consistent within a body of literature. In other words, coordinate-based meta-analyses are blind to effect size magnitude, but direction is tied to the analysis.^55^ Although the coordinate-based meta-analytic approach is limited to studies that report suprathreshold coordinates for a given contrast, this approach will continue to be the most common method for conducting valid meta-analyses for neuroimaging studies until it becomes standard practice to release unthresholded statistical maps.

### Statistical Analysis

All analyses were performed in Montreal Neurological Institute (MNI) space. Coordinates reported in Talairach space were converted to MNI using the “icbm2tal” transformation.^64^ We assessed statistical significance and corrected for multiple comparisons using the permutation-based approach (N = 1000) recommended by the developers of GingerALE.^51,56^ This approach utilized a cluster-forming threshold of *p* < 0.001 (uncorrected) and maintained a cluster-level family-wise error rate of 5%.^56^ Additionally, we took steps to minimize within-group effects on the meta-analyses.^63^ If a study reported more than one contrast (often referred to as an “experiment” in coordinate-based meta-analyses), the contrasts examining similar processes were pooled together to avoid double counting the same participants in a meta-analysis. For example, when a study reported between-group effects in response to $1.50 and $5 rewards relative to neutral or loss conditions, the coordinates derived from the two contrasts were coded as a single reward experiment.

To capture anatomical variation between individual human brains,^65^ we show probabilistic anatomical labels for the locations of the maximum ALE values using the Harvard–Oxford cortical and subcortical atlases.^66^ For transparency, all of the statistical maps (thresholded and unthresholded) derived from the meta-analyses are publicly available on NeuroVault at https://neurovault.org/collections/3884. Readers are free to access these maps and define brain regions using their own labels. Other study materials are available on Open Science Framework at https://osf.io/sjb4d.

## Results

As shown in Figure 1, the systematic literature search identified a total of 41 neuroimaging studies that met the inclusion criteria, yielding 4 coordinate-based meta-analyses with at least 17 independent experiments. Supplementary Tables 2 and 3 show the characteristics of the included studies and their samples.

**Table 2.** Peak Coordinates of Group Differences in Neural Responses to Punishment

| Contrast | Cluster Size (mm<sup>3</sup>) | Probabilistic Anatomical Label | x | y | z |
| --- | --- | --- | --- | --- | --- |
| MDD > HC | 1104 | Amygdala (85.4%) | -26 | -8 | -14 |
|  |  | Amygdala (61.4%) | -16 | -2 | -18 |

Abbreviations: MDD, major depressive disorder; HC, healthy controls. Coordinates are x,y,z
values of the locations of the maximum activation likelihood estimation (ALE) values in MNI space. Probabilistic labels reflect the probability that a coordinate belongs to a given region derived from the Harvard-Oxford probabilistic atlas. For clarity, we only report labels whose likelihood exceeds 5%.

In the present meta-analytic dataset, for the MDD group, the mean number of participants was 19.9, the mean age was 36.4, the mean percentage of females was 60.9%, and the mean percentage of medication usage was 36.6%. For the HC group, the mean number of participants was 20.1, the mean age was 34.9, and the mean percentage of females was 60.3%. Types of reward or punishment used by the included studies encompass money, points, or voucher (41.5%; 17/41); faces (34.1%; 14/41); pictures (12.2%; 5/41); words, statements, captions, or paragraphs (12.2%; 5/41); and autobiographical memory (4.9%; 2/41). Both reward and punishment contrasts were reported in 41.5% (17/41) of studies; 29.3% (12/41) of studies reported punishment contrasts only; and 26.8% (11/41) of studies reported reward contrasts only.

We first synthesized results of 22 studies reporting less activity in response to reward in people with MDD than HC (i.e. HC > MDD for reward > punishment or neutral stimuli and/or neutral stimuli > punishment). As expected, results indicated that these studies reliably reported less activation in a single cluster extending bilaterally across the VS and including part of the subcallosal cortex in MDD (Table 1; Figure 2A).

**Figure 2.**
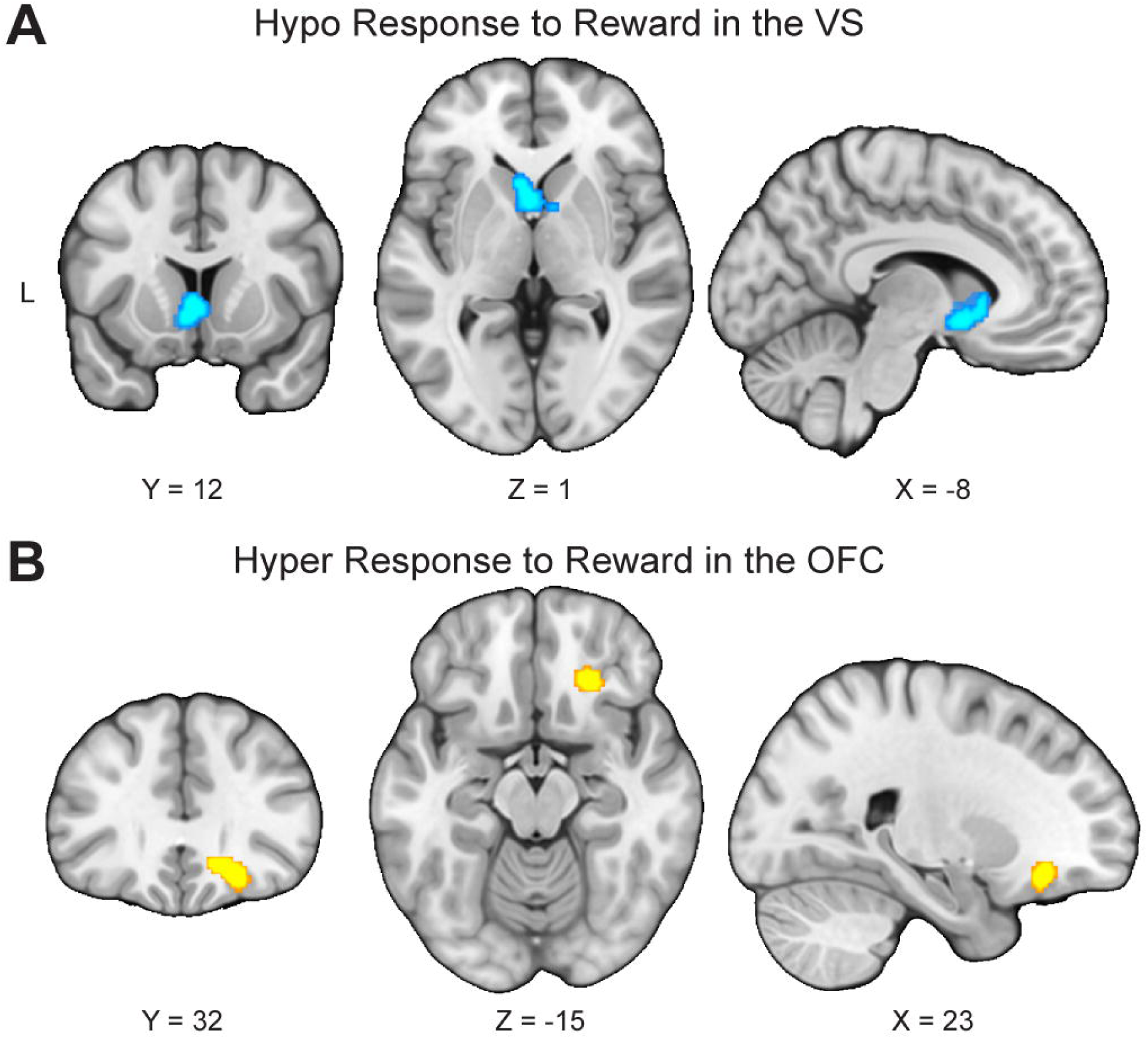
Opposing abnormalities in the reward circuit in response to reward in major depressive disorder. **(A)** To examine regions that consistently showed blunted response to reward, 22 studies reporting less activity in response to reward in people with major depressive disorder (MDD) than healthy controls (HC) were synthesized. Results indicated that these studies reliably report less activation in the ventral striatum (VS) in MDD. **(B)** To identify regions that consistently showed hyper-responses to reward, 18 studies reporting greater activity in response to reward in people with MDD than HC were meta-analyzed. Results indicated that these studies reliably report greater activation in the right orbitofrontal cortex (OFC) in MDD.

In addition to examining which regions consistently showed hypo-responses to reward, we also examined which, if any, brain regions showed consistent hyper-responses to rewarding stimuli. We aggregated results of 18 studies reporting greater activity in response to reward in people with MDD than HC (i.e. MDD > HC for reward > punishment or neutral stimuli and/or neutral stimuli > punishment). Importantly, results indicated that these studies reliably reported greater activation in the right OFC (in a region located in between the medial OFC and the lateral OFC) in MDD (Table 1; Figure 2B).

We conducted sensitivity analyses to examine whether excluding studies that used neutral stimuli > punishment as a reward contrast would affect the main results related to reward responses in MDD. After excluding the two experiments of neutral stimuli > punishment,^38,67^ the results remained the same: We found significant convergence among experiments reporting blunted responses for reward in MDD relative to HC in the VS, as well as significant convergence among experiments reporting elevated responses for reward in MDD relative to HC in the OFC (Supplementary Table 4).

We also conducted exploratory analyses to examine which brain regions consistently show aberrant responses to punishment in MDD relative to HC. First, we meta-analyzed 24 studies reporting greater activity in response to punishment in people with MDD than HC (i.e. MDD > HC for punishment > reward or neutral stimuli and/or neutral stimuli > reward). The results indicated that these studies reliably reported greater activation in the left sublenticular extended amygdala in MDD (Table 2; Figure 3). Second, we synthesized 17 studies reporting less activity in response to punishment in people with MDD than HC (i.e. HC > MDD for punishment > reward or neutral stimuli and/or neutral stimuli > reward). The results indicated that these studies did not report consistent activation patterns. Together, these results suggest that relative to HC, people with MDD exhibited hyper-responses in the left sublenticular extended amygdala during processing of punishment-relevant stimuli.

**Figure 3.**
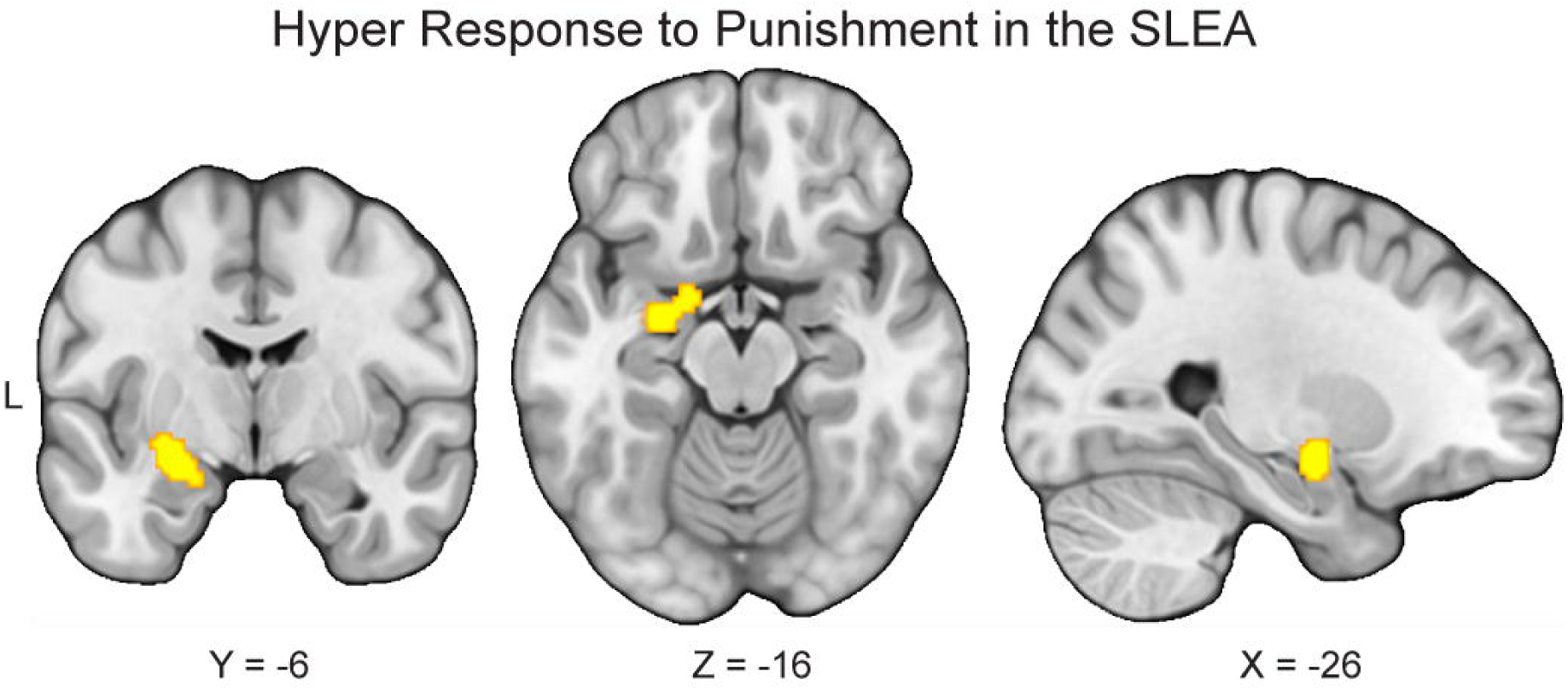
Hyper-responses to punishment in the sublenticular extended amygdala in major depressive disorder. To conduct exploratory analyses to examine which brain regions consistently show elevated response to punishment in major depressive disorder (MDD) relative to healthy controls (HC), we meta-analyzed 24 studies reporting greater activity in response to punishment in people with MDD than HC. The results indicated that these studies reliably report greater activation in the left sublenticular extended amygdala (SLEA) in MDD.

## Discussion

A growing number of researchers have used neuroimaging methods to enhance our understanding of the underlying pathophysiology of MDD.^68–70^ Many of these studies have shown that patients with MDD exhibit blunted responses in the VS, but more disparate patterns of responses in other brain areas. Therefore, it remains unclear what brain regions, other than the VS, are most consistently implicated in people with MDD, particularly during reward processing (see Supplementary Table 1 and Supplementary Figure 1). To address this issue, we performed a coordinate-based meta-analysis of 41 neuroimaging studies containing reward-related responses from a total of 794 patients with MDD and 803 HC. Our meta-analytic findings confirm that reward responses within the VS are consistently blunted in MDD relative to HC across studies. In contrast, we find that reward responses within the OFC are consistently elevated in MDD. Contrary to the common notion that MDD is characterized by blunted responses to reward, these findings suggest that MDD may be characterized by both hypo- and hyper-responses to reward at the neural level and highlight the need for a more fine-tuned understanding of the various components of reward processing in MDD.

Although our blunted striatal findings are in line with previous meta-analytic work documenting reward processing abnormalities in MDD,^34,35,50^ we emphasize that our work differs in two key ways. First, our results implicate highly specific—yet distinct— abnormalities in the reward circuit, with hypo-responses to reward in the VS and hyper-responses to reward in the OFC. In sharp contrast, previous meta-analyses have generally reported distributed patterns of abnormalities, with little anatomical agreement across studies (see Supplementary Table 1 and Supplementary Figure 1). Second, to minimize bias, our study employed more stringent analysis methods than prior studies in this area, following recommendations by new guidelines.^52,55^ For example, instead of using the FDR approach which has been shown to be inadequate in controlling the false positives among clusters in neuroimaging meta-analyses,^53,54^ we corrected for multiple comparisons using the permutation-based approach. We also excluded ROI- or SVC-based studies and only included whole-brain studies that used an active control condition and independent samples. In addition, we only conducted a meta-analysis when there were at least 17 eligible experiments to restrict excessive contribution of any particular studies to cluster-level thresholding.^56^ We speculate that the enhanced rigor and methods of our study contributed to our ability to identify highly circumscribed and distinct abnormalities in the reward circuit.

A prior meta-analysis using similarly rigorous methods revealed no significant convergence of findings among neuroimaging studies comparing MDD and HC.^52^ Nevertheless, we note that the previous meta-analysis differed from the current meta-analysis in at least four key ways. First, whereas the previous meta-analysis focused on emotional or cognitive processing, the current meta-analysis focused solely on reward processing. Second, the previous meta-analysis excluded participants younger than 18 years old; in contrast, the current meta-analysis included participants of all ages, boosting its power and ability to generalize its findings to MDD across ages. Finally, the previous meta-analysis excluded MDD participants in remission, whereas the current meta-analysis included them, leading to the question of whether reward processing dysfunction is not simply a state, but a trait of MDD. This issue needs to be addressed explicitly when there are enough studies available to compare current to remitted depression. Our ability to identify significant convergence highlights the significance of reward processing dysfunction in MDD and might indicate the literature on reward processing in MDD is more homogeneous than that on emotional or cognitive processing in MDD.

In our view, the most important current finding is that studies consistently report that people with MDD exhibit hyper-responses to reward in an OFC region located between the medial OFC and the lateral OFC.^71^ Exposure to rewards (e.g., money and pleasant sights) evokes activity in the OFC, which has been associated with the computation of reward value^18–24^ as well as the representation of current stimulus-outcome associations.^72,73^ For example, whereas the lateral OFC is responsive to stimuli of both positive and negative valence, the medial OFC is particularly responsive to cues of positive valence.^74–76^ Therefore, given that MDD is traditionally linked to blunted response to reward or reduced capacity to experience pleasure,^9^ the finding of hyperactivity of the OFC in response to reward in MDD may seem paradoxical. One interpretation would be that MDD is at least partly characterized by hyper-responses to reward, which fits with a set of experimental studies reporting that individuals with severe MDD found dextroamphetamine to be more rewarding than did controls.^77–79^

Alternatively, OFC hyperactivity may reflect enhanced inhibitory control over subcortical regions underlying reward-related behavior, leading to anhedonia. Optogenetic and neuroimaging studies have revealed that hyperactivity in prefrontal regions (e.g., medial PFC, vmPFC) innervated by glutamatergic neurons may causally inhibit reward-related behavior via suppressing striatal responses to dopamine neurons in the midbrain^7,26^ and increasing connectivity between the medial PFC, lateral OFC, and VS.^7,26^ These findings from animal studies suggest that prefrontal regions (e.g., medial PFC, vmPFC, and lateral OFC) might exert inhibitory control over subcortical regions underlying reward-relevant behavior, potentially explaining the central role of anhedonia in MDD. Nevertheless, it is important to note that given the complexity of the OFC and the inability of the meta-analysis to unravel its varied functions, these interpretations remain speculative.

The extent to which corticostriatal connectivity during reward processing is disrupted in MDD remains an open and important question.^2,80,81^ Previous meta-analyses indicate that at least some people with MDD exhibit dysfunction in resting-state corticostriatal connectivity.^80,81^ The current meta-analytic results will provide a springboard for future studies that seek to develop a full picture of the pathophysiology of MDD and understand the role of dysregulated corticostriatal connectivity in MDD, particularly in the context of reward processing. These endeavors may require empirical assessments of connectivity within the reward circuit using psychophysiological interaction analysis^82–84^ and dynamic causal modeling.^85^ Such approaches have shown promise for revealing specific patterns of task-dependent corticostriatal interactions in samples containing healthy individuals,^86^ clinical populations,^2,87^ or a mix of both.^88^ Nevertheless, a caveat of such approaches is that dysregulated corticostriatal connectivity may involve modulatory regions, such as the midbrain.^89^

In addition to distinct abnormalities within the reward circuit, the current study also found that MDD is associated with hyper-responses in the left sublenticular extended amygdala in response to punishment. The current finding fits with others in suggesting that amygdala hyperactivation is linked to the processing of affectively salient, especially punishing, stimuli in MDD, and may underlie negativity bias in depression.^90,91^ It is also in agreement with a meta-analysis indicating increased activation in the amygdala in response to negative stimuli in MDD relative to HC^68^ and a series of studies indicating that the amygdala may be a key brain region implicated in the pathophysiology of depression.^92,93^ Interestingly, longitudinal studies have reported that amygdala reactivity, potentially in combination with life stress, prospectively predicts internalizing (e.g., depressive and anxiety) symptoms,^94,95^ highlighting the importance of amygdala reactivity in the course of depression.

Even though the current meta-analysis reveals circumscribed patterns of abnormal responses to reward and punishment, we note that the findings should be interpreted in the context of their limitations. First, heterogeneity across studies may have added noise to the current analyses and restricted its capacity for detecting true effects. Specifically, due to the limited number of studies, the current analyses collapsed across different reward processes (e.g., anticipation and outcome), reward modalities (e.g., monetary and social), and specific contrasts that would help isolate and differentiate neural responses to salience and valence.^58^ The current analyses also collapsed across different mood states, psychotropic medication usage, ages, and comorbidities. In doing so, important differences in brain activation may be obscured and more specific questions related to brain activation—particularly questions related to neural representations of valence or salience—cannot be addressed in the current work. Future studies should examine how these factors may affect reward processing in MDD. Nevertheless, we highlight that the convergence of findings despite the heterogeneity of the included studies is striking and suggests that the current findings may reflect central abnormalities of MDD.

Second, some included studies had relatively small sample sizes and reported coordinates that were not corrected for multiple comparisons, which may lead to biased results.^45^ Future work should follow the most updated guidelines for best practices in the field to avoid generating biased findings.^96^ Third, most of the included studies only recruited adults with acute major depression. More studies on other ages (e.g., pre-adolescents, adolescents) and mood states (e.g., remission) are needed. Fourth, we note that the search criteria were designed to focus on studies on reward and might not have identified some studies on punishment. Therefore, the analyses and results in relation to punishment are exploratory in nature and should be interpreted with caution. Fifth, the ALE method, by nature, cannot incorporate null results.^55^ As a result, the current findings could be confounded by publication bias. Sixth, coordinate-based meta-analyses inherently are blind to effect sizes. Therefore, no claims about the magnitude of the effects observed in the current study could be made. Finally, some patients in the included studies were medicated. The normalizing effects of treatment could obscure differences between MDD and HC, increasing the probability of type II errors.^97^

Notwithstanding these caveats, the current meta-analysis shows that MDD is consistently associated with opposing abnormalities in the reward circuit in response to reward: hypo-response in the VS and hyper-response in the OFC. The current meta-analytic results therefore suggest that MDD is associated with both hypo and hyper responses to reward. MDD may stem, in part, from dysregulated connectivity between the VS and the OFC. We believe the current findings will help lay a foundation towards developing a more refined understanding and treatment of MDD and its comorbid psychiatric disorders, particularly ones that involve abnormal reward processing.^98^ For example, a more refined understanding of the abnormalities in the reward circuitry in MDD may help distinguish it from other disorders exhibiting reward processing abnormalities, such as bipolar disorder, schizophrenia, and substance use disorder. Moreover, although reinforcement learning models, such as actor-critic models and prediction error models, have been utilized to understand the pathophysiology of several psychiatric disorders (e.g., schizophrenia and addiction), research on their application on MDD has been scant.^99^ The current results help delineate specific abnormalities within the reward circuit and supply a foundation for refining connectivity-based and computational models of MDD. Finally, given that previous treatment targets for deep brain stimulation for treatment-resistant depression have yielded mixed results,^32^ the portion of OFC implicated by the current results could be a promising treatment target.

## Acknowledgements

This work was supported, in part, by grants from the National Institutes of Health (R21-MH113917 and R03-DA046733 to DVS and R01-MH077908 to LBA).

