## Supplemental Information for "Meta-analysis of Reward Processing in Major Depressive Disorder Reveals Distinct Abnormalities within the Reward Circuit"

***Supplementary Information***

**Supplementary Table 1**

*Comparison of Findings on Reward Responses (i.e., Reward > Punishment/Neutral) in Previous Meta-analyses*

| Brain Region | MNI Coordinates | | |
| --- | --- | --- | --- |
|  | x | y | z |
| *Groenewold et al.* ^19^ | | | |
| *MDD > HC* |  |  |  |
| Lingual Gyrus | 26 | -92 | -14 |
| Olfactorius Cortex | 4 | 22 | -14 |
| Middle Orbitofrontal | 2 | 26 | -14 |
| Rectus | 2 | 30 | -24 |
| Middle Orbitofrontal | 0 | 26 | -12 |
| Rectus | 0 | 24 | -24 |
| *HC > MDD* |  |  |  |
| Cerebellum | -16 | -74 | -28 |
| Lingual Gyrus | -18 | -62 | -6 |
| Fusiform Gyrus | -22 | -74 | -14 |
| Inferior Occipital Gyrus | -30 | -80 | -12 |
| Rolandic Operculum | -40 | -24 | 20 |
| Insula | -36 | -24 | 22 |
| Superior Temporal Gyrus | -40 | -36 | 12 |
| Heschl Gyrus | -46 | -16 | 12 |
| Postcentral Gyrus | -50 | -18 | 18 |
| Supramarginal Gyrus | -50 | -22 | 18 |
| Anterior Cingulate Cortex | -2 | 28 | 16 |
| Anterior Cingulate Cortex | 4 | 32 | 14 |
| Lingual Gyrus | -18 | -62 | -6 |
| Cerebellum | -6 | -58 | -4 |
| Calcarine Sulcus | -20 | -54 | 4 |
| Fusiform Gyrus | -26 | -58 | -12 |
| Precuneus | -20 | -52 | 2 |
| Pallidum | 18 | 0 | -4 |
| Putamen | 28 | -4 | 8 |
| Thalamus | 14 | -8 | 0 |
| Insula | 38 | 10 | -12 |
| Amygdala | 30 | -2 | -12 |
| Caudate | 16 | 26 | 6 |
| Fusiform | 44 | -62 | -20 |
| Crus Cerebellum | 44 | -64 | -20 |
| Brain Region | TAL Coordinates | | |
|  | x | y | z |
| *Zhang et al.* ^20^ | | | |
| *MDD > HC* |  |  |  |
| Cuneus | 4 | -86 | 18 |
| Cuneus | -6 | -86 | 22 |
| Frontal Lobe | 20 | 30 | -6 |
| Middle Frontal Gyrus | 40 | 28 | 38 |
| Superior Frontal Gyrus | -4 | 48 | 32 |
| Fusiform Gyrus | -48 | -74 | -12 |
| Middle Frontal Gyrus | -48 | 14 | 30 |
| Lingual Gyrus | 12 | -52 | 4 |
| Lingual Gyrus | 14 | -54 | 0 |
| *HC > MDD* |  |  |  |
| Caudate | -6 | 18 | 4 |
| Caudate | -8 | -8 | 10 |
| Thalamus | -10 | -12 | 8 |
| Thalamus | -14 | -14 | 16 |
| Caudate | -12 | -4 | 20 |
| Cerebellum | 4 | -36 | -4 |
| Cerebellum | -4 | -42 | 4 |
| Putamen | 14 | 8 | 2 |
| Caudate | 14 | 14 | 10 |
| Anterior Cingulate | -8 | 30 | 10 |
| Insula | 34 | -4 | 16 |
| Cerebellum | -6 | -60 | -20 |
| Brain Region | Coordinates | | |
|  | x | y | z |
| *Keren et al.* ^21^ | | | |
| *HC > MDD* |  |  |  |
| Caudate Body | 12 | 14 | 14 |
| Caudate Head | 6 | 2 | -2 |
| Caudate Body | -8 | -2 | -18 |

Abbreviations: MNI, Montreal Neurological Institute space; MDD, major depressive disorder; HC, healthy controls; TAL, Talairach space. Ventral striatum is the only area implicated in reward processing in MDD relative to HCs across the two previous meta-analyses and the current meta-analysis (see Table 1 for peak coordinates of group differences in neural responses to reward found in the current meta-analysis).

**Supplementary Table 2**

*Characteristics of the Study Samples Included in the Meta-Analysis.*

|  |  | MDD Patients | | | | |  |  | Healthy Controls | | |
| --- | --- | --- | --- | --- | --- | --- | --- | --- | --- | --- | --- |
| Study | Diagnostic Criteria | *n* | Age | % Female | % Medicated | Mood States | Comorbidity |  | *n* | Age | % Female |
| Arrondo *et al.* ^22^ | DSM-IV | 24 | 33.1 | 29.2% | 54.2% | D | Exclusion of alcohol or drug dependence. |  | 21 | 34.3 | 23.5% |
| Bremner *et al.* ^23^ | DSM-IV | 18 | 40 | 66.7% | 0.0% | D | Exclusion of organic mental disorders or comorbid psychotic disorders, post-traumatic stress disorder, childhood trauma, alcohol or substance abuse or dependence, or dyslexia. No current or past history of comorbid psychiatric disorders. |  | 9 | 35 | 77.8% |
| Burger *et al.* ^24^ | DSM-IV | 36 | 40.7 | 61.1% | 100.0% | D | Exclusion of substance dependence. Inclusion of PD, agoraphobia, generalized anxiety disorder, social phobia, obsessive compulsive disorder, post-traumatic stress disorder, somatoform disorder, eating disorder, dysthymia, alcohol abuse, and substance abuse. |  | 36 | 41.3 | 52.8% |
| Chantiluke *et al.* ^25^ | DSM-IV | 20 | 16.2 | 50.0% | 0.0% | D | Exclusion of major psychiatric disorders. |  | 21 | 16.3 | 52.4% |
| Chase *et al.* ^26^ | DSM-IV | 40 | 31 | 77.5% | 77.5% | D | No exclusion of psychiatric comorbidities. Inclusion of lifetime comorbid anxiety disorders and substance use disorders. |  | 37 | 33.1 | 67.6% |
| Demenescu *et al.* ^1^ | DSM-IV | 59 | 36.2 | 66.1% | 23.7% | D | Exclusion of axis I disorders, such as psychotic disorder or dementia, current alcohol or substance abuse. |  | 56 | 39.8 | 60.7% |
| Dichter *et al.* ^27^ | DSM-IV | 19 | 23.6 | 78.9% | 0.0% | R | Exclusion of current axis I psychopathology. |  | 19 | 27.9 | 63.2% |
| Elliott *et al.* ^28^ | DSM-IV | 10 | 42.2 | 70.0% | 100.0% | D | Exclusion of current comorbid anxiety disorders, substance abuse or dependence, bipolar disorder, or other psychiatric diagnoses. Inclusion of past history of PD and bulimia. |  | 11 | 37.6 | 72.7% |
| Engelmann *et al*. ^29^ | DSM-IV | 19 | 37.6 | 52.6% | 0.0% | D | Exclusion of lifetime bipolar disorder, psychotic disorder, obsessive-compulsive disorder, tic disorder, eating disorder, cognitive disorder, substance abuse or dependence in the previous 6 months or positive urine drug screen, or clinically significant suicidal ideation. |  | 23 | 33.7 | 60.9% |
| Fournier *et al.* ^30^ | DSM-IV | 26 | 30.6 | 69.0% | 69.2% | D | Exclusion of bipolar disorder, borderline personality disorder, and alcohol/substance use disorder within 2 months before the scan. Inclusion of history of anxiety disorder and substance abuse. |  | 28 | 32.6 | 57.0% |
| Fu *et al.* ^31^ and ^32^ | DSM-IV | 19 | 43.2 | 68.4% | 100.0% | D | Exclusion of current axis I disorder and history of substance abuse within 2 months of study participation. |  | 19 | 42.8 | 57.9% |
| Fu *et al.* ^33^ | DSM-IV | 16 | 40 | 81.3% | 0.0% | D | Exclusion of other axis I disorder, including anxiety disorder or history of substance within 2 months of study participation. |  | 16 | 39.2 | 81.3% |
| Gotlib *et al.* ^34^ | DSM-IV | 18 | 35.2 | 72.2% | 50.0% | D | Exclusion of psychotic ideation, social phobia, PD, mania, or substance abuse in the past 6 months or behavioral indications of possible impaired mental status. |  | 18 | 30.8 | 72.2% |
| Gradin *et al.* ^35^ | DSM-IV | 25 | 25.5 | 68.0% | 0.0% | D | Unspecified |  | 25 | 25.4 | 68.0% |
| Hall *et al.* ^36^ | DSM-IV | 29 | 37.4 | 55.2% | 51.7% | D | Exclusion of history of alcohol or substance abuse. |  | 25 | 37.7 | 55.2% |
| Johnston *et al.* ^37^ | DSM-IV/ ICD-10 | 19 | 50.8 | 78.9% | 85.0% | D | Exclusion of other primary psychiatric disorder and substance misuse. |  | 21 | 46.1 | 71.4% |
| Keedwell *et al.* ^38^ | ICD-10 | 12 | 43 | 66.7% | 66.7% | D | Exclusion of other axis I disorder. |  | 12 | 36 | 66.7% |
| Knutson *et al.* ^39^ | DSM-III-R | 14 | 30.7 | 64.3% | 0.0% | D | Exclusion of other current axis I disorder. |  | 12 | 28.7 | 66.7% |
| Kumari *et al.* ^40^ | DSM-IV | 6 | 47 | 100.0% | Unspecified | D | Unspecified |  | 6 | 44 | 100.0% |
| Laurent *et al.* ^41^ | DSM-IV | 11 | 24.1 (whole sample) | 100.0% | 23.1% | D | No exclusion of psychiatric comorbidities. Inclusion of past substance abuse/dependence, anxiety disorders, and eating disorder. |  | 11 | 24.1 (whole sample) | 100.0% |
| Liu *et al*. ^42^ | DSM-IV | 21 | 30.7 | 57.1% | 0.0% | D | Exclusion of axis I disorders (other than anxiety) and psychotic features and lifetime substance abuse or dependence. |  | 17 | 28.3 | 58.8% |
| Murrough *et al.* ^43^ | DSM-IV | 20 | 38.1 | 44.4% | 0.0% | D | Exclusion of lifetime history of psychotic illness or bipolar disorder and current alcohol or substance abuse. |  | 20 | 35 | 45.0% |
| Pizzagalli *et al.* ^44^ | DSM-IV | 30 | 43.2 | 50.0% | 0.0% | D | Exclusion of other axis I disorder except for anxiety disorders. |  | 31 | 38.8 | 41.9% |
| Remijnse *et al.* ^45^ | DSM-IV | 20 | 35 | 40.0% | 0.0% | D | Exclusion of current alcohol or substance abuse at the time of study participation. Inclusion of social anxiety disorder, generalized anxiety disorder, PD without agoraphobia, PD, and cannabis abuse in early and sustained full remission. |  | 27 | 32 | 70.4% |
| Rizvi *et al.* ^46^ | DSM-IV | 21 | 38.9 | 66.7% | 0.0% | D | Exclusion of other primary axis I disorder, lifetime history of hypomania/mania, psychosis, obsessive compulsive disorder, or eating disorder, and substance abuse or dependence (except nicotine or caffeine) within the last 3 months. |  | 18 | 36.2 | 66.7% |
| Rosenblau *et al.* ^47^ | DSM-IV | 12 | 43.5 | 41.7% | 0.0% | D | Exclusion of other axis I or II disorders. |  | 12 | 45.8 | 41.7% |
| Scheuerecker *et al.* ^48^ | DSM-IV | 13 | 37.9 | 23.1% | 0.0% | D | Exclusion of past alcohol or substance abuse, other mental illnesses, and personality disorders. |  | 15 | 35.5 | 33.3% |
| Schiller *et al.* ^49^ | DSM-IV | 19 | 23.6 | 78.9% | 0.0% | R | Exclusion of current axis I psychopathology. |  | 19 | 27.9 | 63.2% |
| Segarra *et al.* ^50^ | DSM-IV | 24 | 33.1 | 29.2% | 54.0% | D | Exclusion of dependence on alcohol or recreational drugs. |  | 21 | 34.3 | 19.0% |
| Sharp *et al.* ^51^ | DSM-IV | 14 | 13.4 | 100.0% | Unspecified | D | Exclusion of current use of nicotine, illicit drugs, psychotic disorders, bipolar I disorder, learning disabilities, and mental retardation. |  | 19 | 13.7 | 100.0% |
| Smoski *et al.* ^52^ | DSM-IV | 14 | 34.8 | 50.0% | 0.0% | D | Exclusion of current mood disorder, anxiety disorder, psychotic disorder, substance abuse, or active suicidal ideation and history of psychosis or mania. |  | 15 | 30.8 | 60.0% |
| Smoski *et al.* ^53^ | DSM-IV | 9 | 34.4 | Unspecified | 44.4% | D | Inclusion of generalized anxiety disorder and binge eating disorder. |  | 13 | 26.2 | Unspecified |
| Surguladze *et al.* ^54^ | DSM-IV | 16 | 42.3 | 37.5% | 100.0% | D | Exclusion of illicit substance abuse. |  | 14 | 35.1 | 42.9% |
| Surguladze *et al.* ^55^ | DSM-IV | 9 | 42.8 | 44.4% | 100.0% | D | Exclusion of illicit substance abuse and other axis I disorders. |  | 9 | 39.7 | 44.4% |
| Townsend *et al.* ^56^ | DSM-IV | 15 | 45.6 | 40.0% | 0.0% | D | Exclusion of comorbid axis I disorder. |  | 15 | 44.8 | 40.0% |
| Wagner *et al.* ^2^ | DSM-IV | 19 | 39.9 | 55.0% | 100.0% | D | Exclusion of current comorbid axis I disorder and a history of manic episodes. |  | 20 | 34.1 | 60.0% |
| Wang *et al.* ^57^ | DSM-IV | 12 | 69.1 | 58.3% | 91.7% | D | Exclusion of another major psychiatric disorder and alcohol/drug abuse/dependence. Inclusion of generalized anxiety disorder. |  | 20 | 73.1 | 60.0% |
| Young *et al.* ^58^ | DSM-IV-TR | 16 | 37.1 | 87.5% | 0.0% | D | Exclusion of serious suicidal ideation, psychosis, drug/alcohol abuse in the past year and dependence (except for nicotine) in their lifetime. |  | 16 | 37.8 | 87.5% |
| Zhang *et al.* ^59^ | ICD-10 | 21 | 43.8 | 38.1% | 100.0% | D | Exclusion of illicit substance use or substance use disorders. |  | 25 | 39.3 | 36.0% |
| Zhong *et al.* ^60^ | DSM-IV | 29 | 20.5 | 55.2% | 0.0% | D | Exclusion of lifetime substance dependence and substance abuse in the last 6 months. |  | 31 | 20.8 | 51.6% |

Abbreviations: MDD, major depressive disorder; D, depressed; R, remitted; PD, panic disorder.

**Supplementary Table 3**

*Characteristics of the Studies Included in the Meta-analysis*

| Study | fMRI or PET | Design | Space | Paradigm | Correction | Stimuli | Contrast |
| --- | --- | --- | --- | --- | --- | --- | --- |
| Arrondo *et al.* ^22^ | fMRI | Event-related | MNI | Modified monetary incentive delay task | Uncorrected | Money | HC > MDD, Anticipation: Reward > Non-Reward |
| Bremner *et al.* ^23^ | PET | Block | MNI | Verbal declarative memory tasks with neutral paragraph encoding compared to a control condition and sad word pair retrieval compared to a control condition. | Uncorrected at p < .005 | Words and paragraphs | MDD > HC, Outcome: Negative > Neutral  HC > MDD, Outcome: Negative > Neutral |
| Burger *et al.* ^24^ | fMRI | Event-related | MNI | Face matching paradigm | Corrected at p < .05 (TFCE) | Faces | HC > MDD, Outcome: Negative > Neutral  HC > MDD, Outcome: Positive > Neutral |
| Chantiluke *et al.* ^25^ | fMRI | Event-related | TAL | Reward continuous performance task | Uncorrected at p < .005 | Money | MDD > HC, Outcome: Reward > Non-Reward  HC > MDD, Outcome: Reward > Non-Reward |
| Chase *et al.* ^26^ | fMRI | Event-related | MNI | Card guessing paradigm | Voxel-wise corrected at p < .05 and cluster-wise corrected at p < .01 | Money | MDD > HC, Anticipation: Reward > Non-Reward  HC > MDD, Anticipation: Reward > Non-Reward  MDD > HC, Anticipation: Reward Expectancy  HC > MDD, Anticipation: Reward Expectancy  MDD > HC, Outcome: Prediction Error |
| Demenescu *et al.* ^1^ | fMRI | Event-related | MNI | Viewing faces with angry, fearful, sad, happy, and neutral expressions and scrambled faces; rating gender or pressing buttons in conformity with the instruction presented on the screen | Cluster-wise corrected at p < .05 | Faces | MDD > HC, Outcome: Positive > Scrambled Face |
| Dichter *et al.* ^27^ | fMRI | Event-related | MNI | Modified monetary incentive delay task | Uncorrected at p < .005, k ≥ 10 | Money | MDD > HC, Anticipation: Reward > Non-Reward  MDD > HC, Outcome: Reward > Non-Reward  HC > MDD, Outcome: Reward > Non-Reward |
| Elliott *et al.* ^28^ | fMRI | Block | MNI | Affective go/no go task | Uncorrected at p < .001 | Words | MDD > HC, Outcome: Negative > Positive  HC > MDD, Outcome: Positive > Negative |
| Engelmann et al. ^29^ | fMRI | Event-related | MNI | Economic decision-making task | Cluster-wise corrected at p < .05 | Money | MDD > HC, Outcome: Negative > Positive |
| Fournier *et al.* ^30^ | fMRI | Block | MNI | Labeling a color flash superimposed upon neutral faces that gradually morphed into angry, fearful, sad, or happy faces | Uncorrected at p < .001, k > 20 | Faces | MDD > HC, Outcome: Negative > Neutral MDD > HC, Outcome: Positive > Neutral |
| Fu *et al.* ^31^ and ^32^ | fMRI | Event-related | TAL | Indicating the sex of faces morphed to represent low, medium, and high intensities of sadness | Cluster-wise corrected at p < .005 | Faces | MDD > HC, Outcome: Negative (low, medium, and high intensity)  HC > MDD, Outcome: Positive (low, medium, and high intensity) |
| Fu *et al.* ^33^ | fMRI | Event-related | TAL | Indicating the sex of faces morphed to represent low, medium, and high intensities of sadness | Unspecified | Faces | MDD > HC, Outcome: Negative (low, medium, and high intensity)  HC > MDD, Outcome: Negative (low, medium, and high intensity) |
| Gotlib *et al.* ^34^ | fMRI | Block | MNI | Indicating the sex of faces that were fearful, angry, sad, happy, neutral, or scrambled | Uncorrected at p < .001, k > 5 | Faces | MDD > HC, Outcome: Negative > Neutral  HC > MDD, Outcome: Negative > Neutral  MDD > HC, Outcome: Positive > Neutral  HC > MDD, Outcome: Positive > Neutral |
| Gradin *et al.* ^35^ | fMRI | Event-related | MNI | Ultimatum game | Cluster-wise corrected at p < .05 | Money | HC > MDD, Outcome: Increasing fairness (decreasing inequality)  MDD > HC, Outcome: Increasing inequality (decreasing fairness) |
| Hall *et al.* ^36^ | fMRI | Event-related | TAL | Contingency reversal reward paradigm | Voxel-wise corrected at p < .05 | Money | HC > MDD, Outcome: Magnitude of Loss: Large Loss > Small Loss  HC > MDD, Outcome: Magnitude of Reward: Large Reward > Small Reward  MDD > HC, Outcome: Reward Acquisition > Punishment Reversal  HC > MDD, Outcome: Reward Acquisition > Punishment Reversal |
| Johnston *et al.* ^37^ | fMRI | Event-related | MNI | Modified Pessiglione task | Cluster-wise corrected at p < .01 | Voucher | MDD > HC, Outcome: Loss > Non-Loss  HC > MDD, Outcome: Loss > Non-Loss  MDD > HC, Outcome: Reward > Non-Reward  HC > MDD, Outcome: Reward > Non-Reward |
| Keedwell *et al.* ^38^ | fMRI | Block | TAL | Being exposed to happy, sad, or neutral autobiographical memory prompts and facial expressions | Cluster-wise corrected at p < .01 | Autobiographical memory and faces | MDD > HC, Outcome: Negative > Neutral  HC > MDD, Outcome: Negative > Neutral  MDD > HC, Outcome: Positive > Neutral  HC > MDD, Outcome: Positive > Neutral |
| Knutson *et al.* ^39^ | fMRI | Event-related | TAL | Monetary incentive delay task | Uncorrected at p < .05 | Money | MDD > HC, Anticipation: Reward > Non-Reward  HC > MDD, Anticipation: Reward > Non-Reward  HC > MDD, Outcome: Non-Loss > Loss  HC > MDD, Outcome: Reward > Non-Reward |
| Kumari *et al.* ^40^ | fMRI | Block | TAL | Viewing positive or negative pictures with a caption | Cluster-wise corrected at p < .005 | Pictures and captions | HC > MDD, Outcome: Negative > Neutral  MDD > HC, Outcome: Negative > Neutral  HC > MDD, Outcome: Positive > Neutral  MDD > HC, Outcome: Positive > Neutral  HC > MDD, Outcome: Positive > Negative  MDD > HC, Outcome: Positive > Negative |
| Laurent *et al.* ^41^ | fMRI | Event-related | MNI | Seeing own infant vs. other infant distress faces | Cluster-wise corrected at p < .05 | Faces | HC > MDD, Outcome: Very negative > Negative |
| Liu et al. ^42^ | fMRI | Event-related | MNI | Instrumental probabilistic reward- and punishment-based associative learning task | Cluster-wise corrected at p < .05 | Money | MDD > HC, Outcome: Negative > Neutral  MDD > HC, Outcome: Punishment Prediction Errors |
| Murrough *et al.* ^43^ | fMRI | Event-related | MNI | Rating emotional valence of happy, sad, or neutral faces | Cluster-wise corrected at p < .05 | Faces | HC > MDD, Outcome: 100% Positive > Neutral |
| Pizzagalli *et al.* ^44^ | fMRI | Event-related | MNI | Monetary incentive delay task | Uncorrected at p < .005 | Money | MDD > HC, Anticipation: Loss > Non-Loss  HC > MDD, Anticipation: Loss > Non-Loss  MDD > HC, Anticipation: Reward > Non-Reward  HC > MDD, Anticipation: Reward > Non-Reward  MDD > HC, Outcome: Loss > Non-Loss  HC > MDD, Outcome: Loss > Non-Loss  MDD > HC, Outcome: Reward > Non-Reward  HC > MDD, Outcome: Reward > Non-Reward |
| Remijnse *et al.* ^45^ | fMRI | Event-related | MNI | Reversal learning task | Uncorrected p < .001 | Points | MDD > HC, Outcome: Loss > Baseline  HC > MDD, Outcome: Loss > Baseline  MDD > HC, Outcome: Reward > Baseline |
| Rizvi *et al.* ^46^ | fMRI | Blocked | MNI | Viewing IAPS pictures that elicit positive, negative or neutral affective states | Cluster-wise corrected at p < .05 | Pictures | MDD > HC, Outcome: Positive > Neutral  MDD > HC, Outcome: Negative > Neutral |
| Rosenblau *et al.* ^47^ | fMRI | Event-related | MNI | Viewing IAPS pictures that elicit positive, negative or neutral affective states with and without cues indicating their emotional valence | Uncorrected at p < .05 or p < .005 | Pictures | MDD > HC, Anticipation: Negative > Neutral  MDD > HC, Outcome: Negative > Neutral |
| Scheuerecker *et al.* ^48^ | fMRI | Block | MNI | Face matching paradigm | Uncorrected at p < .001 | Faces | MDD > HC, Outcome: Negative > Neutral |
| Schiller *et al.* ^49^ | fMRI | Event-related | MNI | Monetary incentive delay task | Cluster-wise corrected at p < .05 | Money | HC > MDD, Anticipation: Loss > Non-Loss  HC > MDD, Outcome: Loss > Non-Loss |
| Segarra *et al.* ^50^ | fMRI | Event-related | MNI | Simulated slot-machine game | Cluster-wise corrected at p < .05 | Money | HC > MDD, Outcome: Unexpected Reward > Full Miss |
| Sharp *et al.* ^51^ | fMRI | Event-related | TAL | Card guessing paradigm | Uncorrected at p < .005 | Money | HC > MDD, Outcome: Reward > Non-Reward |
| Smoski *et al.* ^53^ | fMRI | Event-related | MNI | Modified monetary incentive delay task | Cluster-wise corrected | Money | MDD > HC, Anticipation: Money > Control  HC > MDD, Anticipation: Money > Control  MDD > HC, Outcome: Non-Win > Control  HC > MDD, Outcome: Non-Win > Control  MDD > HC, Outcome: Winning > Control  HC > MDD, Outcome: Winning > Control  MDD > HC, Selection: Money > Control  HC > MDD, Selection: Money > Control |
| Smoski *et al.* ^52^ | fMRI | Event-related | MNI | Wheel of fortune task | Uncorrected at p < .005, k ≥ 10 | Money | HC > MDD, Anticipation: Reward > Non-Reward  HC > MDD, Outcome: Reward > Non-Reward |
| Surguladze *et al.* ^55^ | fMRI | Event-related | TAL | Indicating the sex of neutral faces and faces morphed to represent mild and high intensities of fear and disgust | Cluster-wise corrected at p < .001 | Faces | HC > MDD, Outcome: Increasing intensities of happy faces  MDD > HC, Outcome: Increasing intensities of sad faces |
| Surguladze *et al.* ^54^ | fMRI | Event-related | TAL | Indicating the sex of neutral faces and faces morphed to represent mild and high intensities of sadness and happiness | Cluster-wise corrected at p < .001 | Faces | MDD > HC, Outcome: Differential response to 100% disgust  HC > MDD, Outcome: Differential response to 50% fear |
| Townsend *et al.* ^56^ | fMRI | Block | MNI | Face matching paradigm | Cluster-wise corrected at p < .05 | Faces | HC > MDD, Outcome: Negative > Neutral |
| Wagner *et al.* ^2^ | fMRI | Event-related | MNI | Self-referential processing task | Cluster-wise corrected at p < .05 | Statements | MDD > HC, Outcome: Neutral > Negative  MDD > HC, Outcome: Neutral > Positive |
| Wang *et al.* ^57^ | fMRI | Event-related | MNI | Emotional oddball task | Uncorrected at p < .001, k = 5 | Pictures | MDD > HC, Outcome: Negative > Neutral |
| Young *et al.* ^58^ | fMRI | Event-related | TAL | Autobiographical memory task | Cluster-wise corrected at p < .05, k > 30 | Words and autobiographical memories | HC > MDD, Outcome: Very Positive > Positive  HC > MDD, Outcome: Very Negative > Negative  MDD > HC, Outcome: Very Negative > Negative |
| Zhang *et al.* ^59^ | fMRI | Event-related | MNI | Viewing IAPS positive, neutral, and negative pictures with or without valence cues | Cluster-wise corrected at p < .05, k > 157 | Pictures | MDD > HC, Outcome: Reward > Non-Reward |
| Zhong *et al.* ^60^ | fMRI | Block | MNI | Face matching paradigm | Uncorrected at p < .005, k =8 | Faces | MDD > HC, Outcome: Negative > Neutral  HC > MDD, Outcome: Negative > Neutral |

Abbreviations: fMRI, functional magnetic resonance imaging; PET, positron emission tomography; MNI, Montreal Neurological Institute space; SVC, small volume correction; MDD, major depressive disorder; HC, healthy controls; TFCE, threshold-free cluster enhancement; TAL, Talairach space; VS, ventral striatum; dACC, dorsal anterior cingulate cortex; rACC, rostral anterior cingulate cortex; ACC, anterior cingulate cortex; mPFC, medial prefrontal cortex; mOFC, medial orbitofrontal cortex; IAPS, International Affective Picture System.

**Supplementary Table 4**

*Peak Coordinates of Group Differences in Neural Responses to Reward (Excluding Neutral Stimuli > Punishment)*

| Contrast | Cluster Size (mm^3^) | Probabilistic Anatomical Label | x | y | z |
| --- | --- | --- | --- | --- | --- |
| MDD > HC | 968 | Frontal Orbital Cortex (26%), Frontal Pole (13%) | 20 | 32 | -12 |
| HC > MDD | 1784 | Subcallosal Cortex (14%) | -2 | 8 | -4 |
|  |  | Caudate (32.1%),  Accumbens (11.1%) | 8 | 6 | -2 |

Abbreviations: MDD, major depressive disorder; HC, healthy controls. Coordinates are x,y,z values of the locations of the maximum activation likelihood estimation (ALE) values in MNI space. Probabilistic labels reflect the probability that a coordinate belongs to a given region derived from the Harvard-Oxford probabilistic atlas. For clarity, we only report labels whose likelihood exceeds 5%.

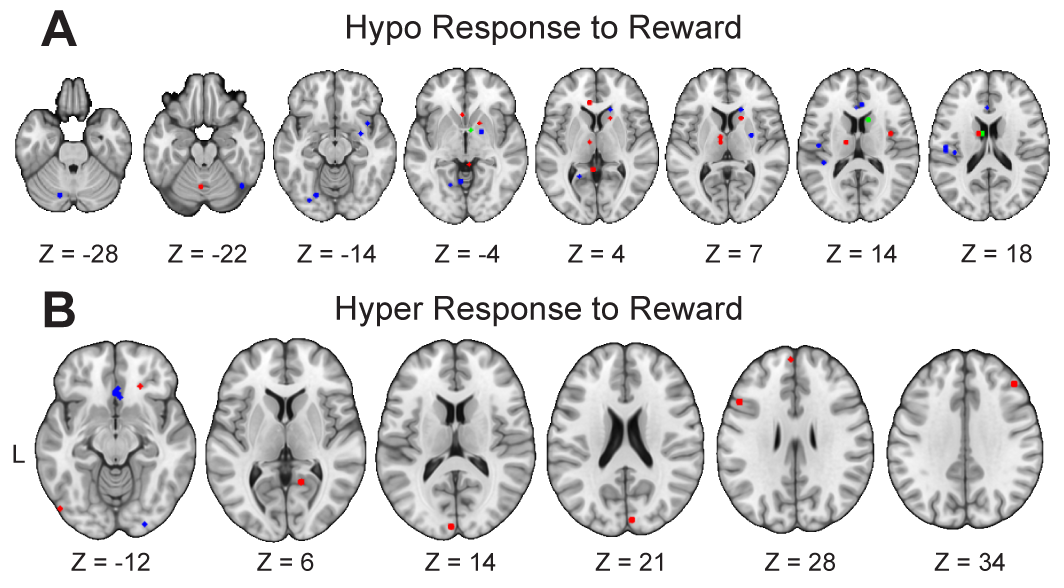

**Supplementary Figure 1**. Illustration of Findings of Previous Meta-analyses on Reward Processing in Unipolar Depression. There is a striking degree of anatomical disagreement across these meta-analyses, with non-overlapping findings all throughout the brain. Blue represents Groenewold et al.^19^ Green represents Keren et al.^21^ Red represents Zhang et al.^20^ **(A)** Previous meta-analyses examining convergence among studies reporting hypo-responses to reward include Groenewold et al.,^19^ Keren et al.,^21^ and Zhang et al.^20^ **(B)** Previous meta-analyses examining convergence among studies reporting hyper-responses to reward include Groenewold et al.^19^ and Zhang et al.^20^

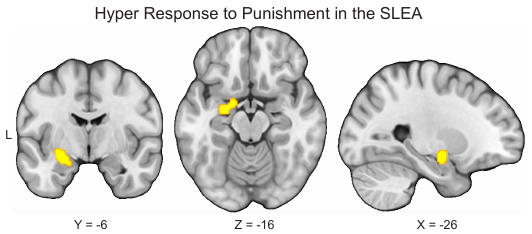

**Supplementary Figure 2**. Hyper-responses to punishment in the sublenticular extended amygdala (SLEA) in major depressive disorder (MDD). To conduct exploratory analyses to examine which brain regions consistently show elevated response to punishment in MDD relative to healthy controls (HCs), we meta-analyzed 24 studies reporting greater activity in response to punishment in people with MDD than HCs. Our results indicated that these studies reliably report greater activation in the left SLEA in MDD.
